# *SplAdder*: Identification, quantification and testing of alternative splicing events from RNA-Seq data

**DOI:** 10.1101/017095

**Authors:** André Kahles, Cheng Soon Ong, Yi Zhong, Gunnar Rätsch

## Abstract

**Motivation**: Understanding the occurrence and regulation of alternative splicing (AS) is a key task towards explaining the regulatory processes that shape the complex transcriptomes of higher eukaryotes. With the advent of high-throughput sequencing of RNA (RNA-Seq), the diversity of AS transcripts could be measured at an unprecedented depth. Although the catalog of known AS events has grown ever since, novel transcripts are commonly observed when working with less well annotated organisms, in the context of disease, or within large populations. Whereas an identification of complete transcripts is technically challenging and computationally expensive, focusing on single splicing events as a proxy for transcriptome characteristics is fruitful and sufficient for a wide range of analyses.

**Results**: We present *SplAdder*, an alternative splicing toolbox, that takes RNA-Seq alignments and an annotation file as input to *i*) augment the annotation based on RNA-Seq evidence, *ii*) identify alternative splicing events present in the augmented annotation graph, *iii*) quantify and confirm these events based on the RNA-Seq data, and *iv*) test for significant quantitative differences between samples. Thereby, our main focus lies on performance, accuracy and usability.

**Availability**: Source code and documentation are available for download at http://github.com/ratschlab/spladder. Example data, introductory information and a small tutorial are accessible via http://bioweb.me/spladder.

## 1 Introduction

Alternative splicing (AS) is an mRNA processing mechanism that cuts and re-joins maturing mRNA in a highly regulated manner, thereby increasing transcriptome complexity. Depending on the organism, up to 95% of expressed genes are transcribed into multiple transcript variants (Pan *et al*., 2008; Wang *et al*., 2008), where various transcripts with differing exon composition can arise from the same gene locus. (Throughout this text, we will use the term *transcript* to identify a variant of a gene that was generated through transcriptional processing.) Although these transcripts might never coexist at the same time and place, each one of them can be essential for cell differentiation, development or play an important role within signaling processes (Kornblihtt *et al*., 2013). Thus, the two major challenges in computational transcriptome analysis are complexity and completeness. In *SplAdder*, we leverage evidence from RNA-Seq data to compute a more complete representation of the splicing diversity within a sample and tackle the complexity with a reduction to alternative splicing events instead of full transcripts. We provide open source implementations for *SplAdder* in MATLAB and Python that contain all features described below and produce the same results. However, future development will focus on the Python implementation for reasons of accessibility. All inputs follow the standardized formats for alignments and annotation such as BAM and GFF. For complete examples, use cases and information regarding the user interface, we provide a supplementary website. User documentation is available in the wiki section of the source code repository.

In Section 2 we will give a brief overview on related approaches that also focus on the analysis and quantification of alternative splicing based on RNA-Seq data. Our main focus will be on methods that are able to characterize alternative splicing events. In the subsequent Section 3, we give an outline of the *SplAdder* methodology and the algorithmic details of its main compute phases. To show how *SplAdder* compares to other strategies for RNA-Seq based alternative splicing analysis, we have compiled a set of different evaluations and comparisons to existing methods. Our experimental design will be described in Section 4 and the main results are discussed in Section 5. Lastly, Section 6 summarizes this work.

## 2 Related Work

Prior to the advent of high throughput RNA-Seq, methods based on expressed sequence tags (ESTs) were developed to elucidate the complex patterns of alternative splicing in higher organisms (Modrek and Lee, 2002). Although designed for a much lower data throughput, the algorithmic ideas presented for ESTs have had a strong influence to the field in the following years. One central idea is the representation of splicing variation at a gene locus as a graph that encodes exon segments as nodes and the intron segments as connecting edges (Heber *et al*., 2002; Eichner *et al*., 2011; Kianianmomeni *et al*., 2014). Similar to *SplAdder*, numerous tools are based on such splicing graph representations; however, none of the existing approaches combines all aspects of the *SplAdder* workflow: the augmentation of existing annotation information, the detection and quantification of alternative splicing events, differential testing of events between two given sets of samples and detailed visualization of the splicing variation. There exist several approaches that cover at least a subset of the steps in the *SplAdder* pipeline. The most notable ones are *JuncBase* (Brooks *et al*., 2011), *rMATS* (Shen *et al*., 2014) and SpliceGrapher (Rogers *et al*., 2012). *JuncBase* utilizes third party prediction tools such as *Cufflinks* (Trapnell *et al*., 2010) to allow for the detection of novel exon nodes in the splicing graph. It then extracts and quantifies splicing events of the most common AS types and reports them in a custom format. Further, *JuncBase* provides basic differential analyses and basic visualizations of the test results. However, the pipeline consists of 10 different steps, including building a *Cufflinks* output based database, which is quite laborious to generate, has a long running-time and is thus not ideal for larger scale studies. *SpliceGrapher* directly integrates information from RNA-Seq or EST data into a splicing graph and can display splicing events in the graph visualizations. Unfortunately, it does not provide an easy method to explicitly generate and quantify alternative splicing events and does not allow for differential analysis. *rMATS* focuses on the differential analysis of splicing between RNA-Seq samples. It can detect the most common AS events from either RNA-Seq alignments or from a set of reads by applying a third party mapping algorithm. Based on the RNA-Seq evidence, it will also fill in some missing information to call events not present in the provided annotation but has a limited capacity to do so.

Other methods, such as *Scripture* (Guttman *et al*., 2010), *Cufflinks* (Trapnell *et al*., 2010) or *MISO* (Katz *et al*., 2010) also use graphs internally and allow for novel splice variants based on RNA-Seq evidence but focus on the prediction of full transcripts instead of single events. These tools aim to solve a much harder problem and thereby miss potential local variability for AS studies. These tools are also computationally more expensive, limiting their applicability in the context of thousands of samples. Another popular tool that is focused on the extraction of alternative splicing events from a given annotated locus is the *Astalavista* toolbox (Foissac and Sammeth, 2007). Although many splicing events are covered in the detection phase, the tool relies on a complete annotation as input and does not provide any quantification values for the events However, the authors introduce a logical representation of splice events (the splicing code) that we will utilize later on. The software *SpliceTrap* (Wu *et al*., 2011) is able to generate quantification values for the most common AS types, but recognizes much fewer transcripts than *Astalavista*. For both tools no novel splice variants are considered.

In our evaluation on simulated data, we will show that *SplAdder* is more accurate in detecting novel events and shows better performance in differential analysis than any of the tested competitors. We have chosen to compare *SplAdder* against *JuncBase*, *rMATS* and SpliceGrapher as these methods are closest to the presented *SplAdder* pipeline. We discuss further details regarding the comparisons in Section 4 and Suppl. Section D.

## 3 Approach

The *SplAdder* algorithm consists of multiple steps that convert a given annotation into a splicing graph, enrich that graph with splicing evidence from RNA-Seq samples, identify splicing events from the augmented graph and use the given RNA-Seq data to quantify the single events (Figure 1). Optionally, the quantifications can then be used for differential analysis. We find this distinction important, as differential analysis between samples is only one of many possible applications of AS event phenotypes. Other examples may include generating of sample specific splicing profiles or using AS phenotypes in genome-wide association studies.

**Figure 1:**
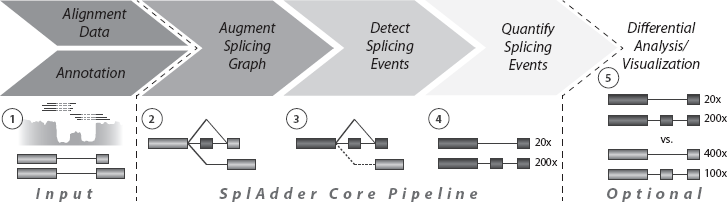
*SplAdder* Analysis Flowchart The main steps of the *SplAdder* workflow consist of (1) integrating annotation information and RNA-Seq data, (2) generating an augmented splicing graph from the integrated data, (3) extraction of splicing events from that graph, (4) quantifying the extracted events, and optionally (5) the differential analysis between samples and producing visualizations.

### 3.1 Preliminaries

Here, we will introduce our notation and make definitions that will be used throughout the following descriptions of the algorithm.

**Coordinates** All positions used in the following descriptions are in a genomic coordinate system. We begin by defining the genome 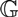 as a string of consecutive positions 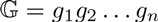. When addressing any range *x* within these positions, *e.g*., to define a gene *x*, we describe this as the pair of the first and the last position of *x*: (*g_x;start_*; *g_x;end_*). When addressing a specific entity *x_i_*, we will write (*g_xi;start_*; *g_xi;end_*). For simplicity, we ignore chromosomes and assume the genome to be one continuous string.

**Representation of Genes as Transcript Graphs** A given gene annotation can be represented as a set of linear directed graphs. Assume gene 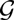 as given, that has *k* different transcripts 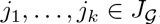, where 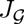 is the set of all transcripts of gene 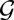. As we consider each gene 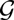 independently, we will omit the index 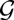 wherever possible in order to keep the notation uncluttered. Each transcript consists of a set of exons that are connected by introns. Each exon can be uniquely identified by its start and end. We thus represent all exons as coordinate pairs of their genomic start and end position:

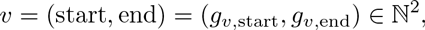

where *g_v,start_* and *g_v,end_* are the first and last position of exon *v* in genomic coordinates, respectively. Although further coordinate information like chromosome and strand are used in the program implementation, we will limit this description to an identification by start and end for simplicity. The exons ofeach transcript *j_i_* can then be represented as a node set *V_i_*: = {*v_i_*,_1_,…, *v_i,mi_*} with 1 ≤ *i* ≤ *k* and *m_i_* as the number of exons in transcript *j_i_*. As transcripts have a direction (the exons within a transcript follow a strict order), we require, that the index of the nodes reflects the order of the exons in the transcript. As no two exons in a transcript overlap by definition, this order is implied by *g_v,start_* and *g_v,end_*. We then define the edge set of transcript *j_i_* as

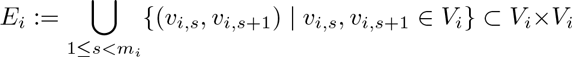

with 1 ≤ *i* ≤ *k*. The pair (*V_i_, E_i_*) forms the directed *transcript graph* of transcript *j_i_*.

**Definition of Splicing Graphs** We define the set of exons occurring in *any* transcript *j_i_* as *V*. As the single exons are uniquely identified by their coordinates, we can write 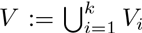. Hence, we define the set of all edges as

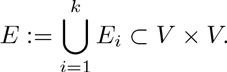

Note, that only already existing edges are merged, preserving the preexisting order of nodes. The pair *G* = (*V, E*) is a directed acyclic graph and is called the *splicing graph* representation of a gene. Figure S-2 illustrates how a set of five transcripts is collapsed into a splicing graph. The key concept is, that when multiple transcripts contain the same exon, this will be represented by a single node in the splicing graph.

We define the *in-degree* and the *out-degree* of a node as the number of its incoming and outgoing edges, respectively. We further define a node to be *start-terminal*, if its in-degree is zero and *end*-terminal if its out-degree is zero. Each transcript can now be represented as a path through the splicing graph, beginning at a start-terminal node and ending at an end-terminal node.

Note, that although the splicing graph representation resolves many redundancies and efficiently stores large numbers of different but mostly overlapping transcripts, this comes at the cost of information loss. Long range dependencies between single exons are not preserved. An example of this is provided in Figure S-2. Although exon T2E1/T3E1 exclusively occurs in transcripts that end in exon T2E3/T3E3, this relationship is lost in the graph, where E2 can connect to both E6 and E7. Our approach is not severely affected by this limitation as we only extract local information about alternative exon- or intron-usage.

**Definition of Segment Graphs** Following the splicing graph definition, two or more nodes in the graph may overlap. Thus, when collecting expression information for each node from a given alignment, the same genomic positions may be queried multiple times. To overcome this inefficiency, we use the concept of breaking down each node into non-overlapping exon segments, similarly used in (Reyes *et al*., 2012; Behr *et al*., 2013).

The same principle that is applied when collapsing different transcripts that share the same exons into a graph structure can also be applied to collapse exon segments that are shared by several nodes of the splicing graph. Following this idea, we divide each exon into non-overlapping segments. Analogous to an exon, a segment is uniquely identified by its genomic coordinate pair and the same order as on exons can be applied: *s* = (*g_s,start_*, *g_s,end_*). We say an exon *v_i_* is *composed* from segments *s_iq_* through *s_i,r_*, if *v_i_* = *s_i_,q*°^s_i,r_^, with *q* < *r* and where .°. denotes the concatenation of segment positions. Thus, the set of all segments can be defined as

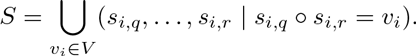

To explicitly define the set of all segments, first we define the set *V_S_* of all node-starts in *V* and the set *V_T_* of all node ends in *V*. The set of all segments *S* can then be defined as

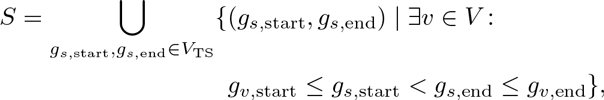

where *V_ST_* = *V_S_* ∪ *V_T_*. The computation of *S* from *V* is straightforward. Let *P* be a sorted array containing all genomic positions that are either start or end positions of an exon in *V*. We denote the *i*-th element of the array as *P[i]*. Let *L_S_* and *L_E_* be two binary label-arrays with the same length as P, where *L_S_[i]* is 1 if *P[i]* is start of an exon in *V* and 0 otherwise. Correspondingly, *L_E_[i]* is 1 if *P[i]* is the end of an exon in *V* and 0 otherwise. Let further *C_S_* and *C_E_* be two arrays with the same length as *P*, where 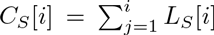 and 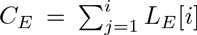 are the cumulative starts and ends up to position *i*. We can then determine the set of all segments as

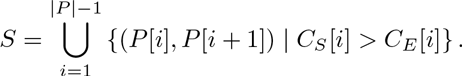

Similar to the definition of the edges for the splicing graph, we define

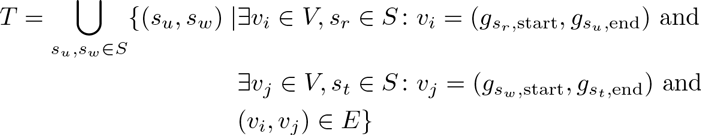

to be the set of segment pairs that are connected by an intron. We then denote the pair *R* = (*S, T*) to be the *segment* graph of a gene. For practical reasons, we store an additional matrix that relates each node in the splicing graph to the segments it is composed of. Supplemental Figure S-5 illustrates the relationship between splicing graph and segment graph.

We will use the splicing graph representation to incorporate new information based on RNA-Seq evidence as well as for the extraction of alternative splicing events. We will use the segment graph representation for event quantification, as this is computationally much more efficient.

### 3.2 Construction of an Augmented Splicing Graph

As a preprocessing step, the input annotation is transformed into the initial splicing graph *G* according to the definitions above, thereby collapsing exons shared by multiple transcripts into single nodes of the graph. In the following, we describe how *G* is transformed into an augmented graph *Ĝ* using information from RNA-Seq data, thereby introducing new nodes and edges. This is an integral part of the *SplAdder* workflow that enables the discovery of novel splicing variation based on RNA-Seq data.

The augmentation of *G* is a four-step algorithm:

1. build initial graph
2. add novel cassette exons
3. add novel intron retentions
4. while novel edges can be added

4.1 insert novel intron edges

When a newly added node shares one boundary with an existing node, the existing edges are inherited by the new node. Following, we will provide a detailed explanation for each step.

Given an RNA-Seq sample and a gene 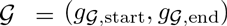, we extract all intron junctions from the alignment that overlap 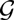 and show sufficient alignment support. Whether an intron junction is sufficiently well supported is based on a set of given confidence criteria (cf. Supplemental Table *C*) We define the list of RNA-Seq intron junctions 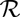 as

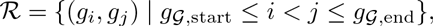

where (*g_i_; g_j_*) describes the intron starting at *g_i_* and ending at *g_j_*. Further, let *v* = (*g_v;start_*; *g_v;end_*), with *v* ∈ *V*, be an existing node in the splicing graph. The augmentation process will transform the existing splicing graph *G* = (*V;E*) into an augmented graph 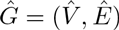. We initialize *Ĝ* with *G*.

**Adding Novel Cassette Exons** In the first augmentation step, new cassette exon structures are added to the splicing graph. For this, the algorithm iterates over all non-overlapping pairs of 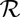. For each pair (*g_i_*_1_; *g_j_*_1_) and (*g_i_*_2_; *g_j_*_2_), two conditions need to be fulfilled. Briey, both intron ends need to be attached to existing exons and the cassette exon must not already exist. Formally, we check for the following conditions:

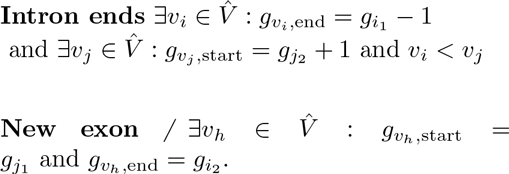

If both conditions are met, a new node *v_n_* = (*g_j_*_1_ + 1; *g_i_*_2_ – 1) is added to the node set 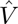 and two new edges (*v_i_; v_n_*) and (*v_n_; v_j_*) are added to *Ê*. Figure S-1, Panel A, schematically describes the addition of a cassette exon. The criteria for adding a cassette exon are listed in Supplemental Table A.

**Adding Novel Intron Retentions** The second augmentation step adds intron retention events to the splicing graph. For each edge (*v_s_*; *v_t_*) ∈ *Ê*, the algorithm decides whether there is enough evidence from the given RNA-Seq sample for expression inside the intron, to consider the intron sequence as retained. Again, heuristic confidence criteria are applied (cf. Supplemental Table B). Briey, the central criteria for adding a new intron retention is the number of sufficiently covered positions within the intron as well as the differences in mean coverage between intronic and exonic part of that region. When sufficient evidence for a retention is found, a new node *v_n_* = (*v_s;start;_ v_t;end_*) is added to 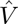. The new node inherits all incoming edges from *v_s_* and all outgoing edges from *v_t_*, thus we get the set of newly added edges

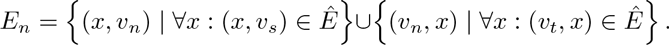

Then, the set of edges is updated with 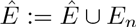. Supplemental Figure S-1, Panel B, illustrates this case.

**Insert Novel Intron Edges** The last augmentation makes once more use of the list of RNA-Seq supported intron junctions 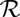 generated during the first step. Based on start and end position of the intron, we can test if any existing nodes start or end at these positions, respectively. We have to distinguish between four different basic cases: 1) neither start nor end coincide with any existing node boundary, 2) the intron-start coincides with an existing node end, 3) the intron end coincides with an existing node-start, 4) both the intron-start coincides with an existing node end and the intron-end coincides with an existing node-start. The four cases and their respective sub-cases are illustrated in Panels C-H of Supplemental Figure S-1. Formal definitions of the different cases are given in Supplemental Section A. As the addition of novel intron edges depends on other possibly novel edges, this addition step is repeated iteratively until no new edges can be added or a pre-defined maximum number of iterations is reached.

**Splicing Graph Pruning** When multiple RNA-Seq samples are available, *SplAdder* allows for an optional filtering step to reduce false positive edges. All edges that are not supported by a given minimum number of RNA-Seq samples will be pruned from the graph. Resulting orphan nodes that were not present in the initial graph will be pruned as well.

### 3.3 Detect and Quantify Alternative Splicing Events

Based on the augmented splicing graph, we extract various classes of AS events as subsets of connected nodes. *SplAdder* currently supports the following event types: exon skip, intron retention, alternativ 3’ and alternative 5’ splice sites, multiple exon skips as well as mutually exclusive exons. Note, that currently alternative transcript starts and ends are not detected, as they are products of alternative tran-scriptional processing rather then results of alternative splicing. Each event is then represented as a “mini-gene” consisting of two splice variants minimally describing the alternatives of the event. Overlapping events that share the same intron coordinates and do only differ in the flanking exon ends are merged into a short common representation. We refer to Supplemental Section B.1 for the formal definitions of all classes of alternative events and a detailed description of the extraction algorithms.

Finally, the event set identified from the splicing graph is quantified using the given read alignment data. For each event, we report the mean coverage of each exon and the number of spliced alignments supporting each intron. Remember, that to speed up the quantification process, the read counting is performed on the segment graph representation defined above. Thus, no exon position needs to be quantified twice.

### 3.4 Differential Analysis

If the set of input samples can be separated into two or more groups representing different conditions, the splice quantifications produced by *SplAdder* can be subjected to differential testing. For this, *SplAdder* provides two basic strategies. The first is to use the *SplAdder* output files that describe event structure and quantification as input to other tools dedicated to analyze differential expression, such as *rDiff* (Drewe *et al*., 2013) or DESeq (Reyes *et al*., 2012). In previous studies, we have generally used the combination of *SplAdder* and *rDiff*. In this case, the mini genes predicted by *SplAdder* are re-quantified by *rDiff* and subjected to a test for differential relative transcript usage.

The second strategy is to directly use the exon-intron junction counts generated by *SplAdder* to apply a differential test. Briefly, we model junction read counts with a negative binomial distribution and employ a generalized linear model (GLM) framework for testing similar to (Love *et al*., 2014). Similar to the previous approach, we use the sample replicate to estimate a mean variance relationship to better account for overdispersion. Details of the GLM based test is provided in Supplemental Section C. This strategy can be run as part of the *SplAdder* pipeline. It directly accesses the event quantifications and is computationally more efficient than the previous hybrid approach. We have included both strategies into our evaluation presented in Section 4.

### 3.5 Visualization

SplAdder also provides means for publication-ready visualization of the RNA-seq read coverage of exon positions and of intron junctions. Visualization allows for effective visual inspection of identified alternative splicing events in light of primary read data. These visualizations provide summarization of multiple samples as well as the comparison of different groups of samples to highlight differential splicing over several replicate groups or conditions. An example is provided in Supplemental Figure S-8.

## 4 Evaluation and Applications

The *SplAdder* approach has been successfully applied in various biological studies on *Arabidopsis thaliana* (Drechsel *et al*., 2013; Gan *et al*., 2011) as well as in the context of large-scale cancer projects with several thousand RNA-seq libraries (Weinstein *et al*., 2013). Here, we have created several sets of simulated data to evaluate *SplAdder*. Simulated data allows for an accurate measure of performance and provides a ground truth for a fair comparison against other existing methods. To allow as little bias as possible towards our own method, we used an external data simulator (Griebel *et al*., 2012). In the following, we describe the generated datasets and which evaluations were performed on them.

### 4.1 Data simulation

**Detection of Novel Events** We have used the FluxSimulator (Griebel *et al*., 2012) toolbox to simulate RNA-Seq data sets of sizes 5 million, 10 million and 20 million reads, covering 1, 000 genes randomly selected from the human GENCODE annotation (v19) (Harrow *et al*., 2012) at various depths. For this analysis, we put our main focus on the sensitive detection of novel alternative splicing events. Thus, we pre-filtered the annotation to genes that had at least two transcripts annotated.

All reads were aligned to the human reference genome using the *STAR* (Dobin *et al*., 2013) as well as the *TopHat2* (Kim *et al*., 2013) aligners to show the applicability of our pipeline in a general context. In both cases, we provided the full reference annotation for index creation. *TopHat2* implements a 2-pass alignment mode per default. As this mode is optional for *STAR*, we ran it with and without 2-pass mode to also get a better understanding of its benefits. In addition to the alignment output, we also transformed the simulated read alignments into BAM format and used it as optimal input for the splice prediction tools, best reflecting ground truth information.

To simulate a realistic scenario of detecting novel AS events based on the provided RNA-Seq alignments only, we provided only a reduced annotation to the tools performing the AS event prediction. This reduced representation contains only the first annotated transcript of a gene, where first is defined as first occurrence in the complete annotation file.

For further details on data set creation and alignment, including all command line parameter settings, we refer to Supplemental Section D.

**Differential Analysis** The simulated data for the analysis of differential testing was taken from the publication of *rDiff* (Drewe *et al*., 2013), a tool for the detection of differentially expressed transcripts from RNA-Seq data. The two datasets consist of 5,785 genes each, where half of the genes shows differential relative transcript expression and the other half does not. The *rDiff* publication gives further details on dataset generation.

### 4.2 Evaluation

**Detection of Novel Events** We used the *Astalavista* toolbox (Foissac and Sammeth, 2007) to extract all annotated alternative splicing events from the set of the randomly chosen 1,000 genes that we used for data simulation. In contrast to the individual prediction tasks, *Astalavista* had access to all annotated transcripts of a gene and thus generated our ground truth set used for evaluation later on. *Astalavista* generates output following a well-defined nomenclature (Guigό Serra *et al*., 2008).

The single AS event predictors were run on the limited annotation containing only the first transcript but had access to the RNA-Seq data generated from the non-constrained annotation set. We then converted the output of all other tools into the well defined *Astalavista* format to allow for an easy comparison. For each of the four AS event types (exon skip, intron retention, alternative 3’ splice site and alternative 5’ splice site), we compared the predictions to the ground truth set and computed precision, recall and F-score metrics.

For this evaluation we considered *JuncBase*, *rMATS*, *SpliceGrapher* and *SplAdder*.

**Event Quantification** Based on the read data simulated for the detection of novel events, we were also able to evaluate the event quantifications provided by the respective approaches. We based all our analyses an percent spliced in (PSI) values, as they are an accepted standard in the community. To generate the ground truth PSI values, we took the relative expression of a transcript for each gene as simulated by *FluxSimulator*. For each alternative splicing event, we computed its PSI value as the ratio between the sum of abundances of transcripts that represented the inclusion (*e.g*., not skipping the exon in an exon skip event) over the sum of abundances of all transcripts containing any of the event exons.

The so generated PSI values were then used as ground truth for comparison of the predicted event quantifications. Only the correctly detected events of each approach could be compared to the ground truth quantifications. We used the Pearson correlation coefficient as a measure of agreement between predicted and true PSI values.

This evaluation was performed for *JuncBase*, *rMATS* and *SplAdder*, as *SpliceGrapher* does not provide quantification values.

**Differential Analysis** The two test sets taken from (Drewe *et al*., 2013) contain 5,785 genes each that either do (2,937) or do not (2,938) show differential transcript usage. One dataset shows small variability and the other large variability, which we will further refer to as the *small* and *large* dataset, respectively. For each dataset, we used the set of differential genes as ground truth and counted a prediction as a true positive if the tool found at least one significant AS event in that gene. From this we generated receiver operating characteristic (ROC) curves with increasing significance cut-offs to evaluate each tool’s performance.

For this analysis we compared only *rMATS*, *JuncBase* and *SplAdder*, as *SpliceGrapher* does provide no differential testing functionality.

## 5 Results

### 5.1 Detection of Novel Events

Based on the three sets of simulated reads and the different alignments performed on these read sets, we evaluated how well the single prediction tools can reconstruct the splicing variability in the sample from read alignments and limited annotation. In comparison to the ground truth dataset generated by using *Astalavista* on the non-restricted annotation file, we computed precision, recall and F-Score metrics for four types of AS events (Figures 2, S-6 and S-7).

**Figure 2:**
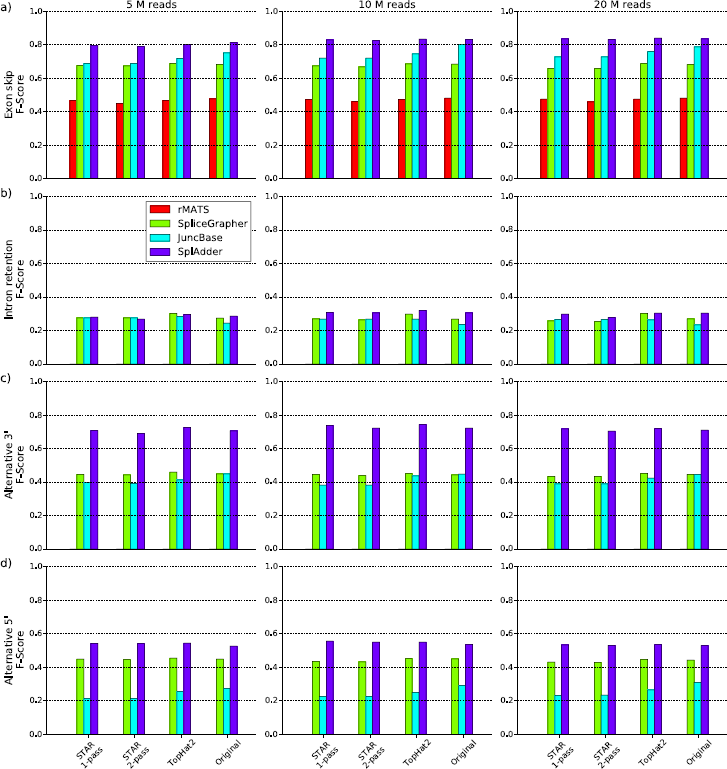
***SplAdder* Evaluation Results** This matrix of bar charts summarizes the evaluation results for the comparison of *rMATS*, *SpliceGrapher*, *JuncBase* and *SplAdder* (see legend) on different sets of simulated RNA-Seq read data. The metric shown here is the F-Score, defined as the harmonic mean of precision and recall. (Plots of the same design with details on precision and recall are provided in Supplemental Figures S-6 and S-7.) The rows of the plot matrix represent four different event types: a) exon skip, b) intron retention, c) alternative 3’ splice site, and d) alternative 5’ splice site. The columns represent different read set sizes (5 million, 10 million, 20 million). The four bar groups represent the different aligners used (from left to right: *STAR* 1-pass, *STAR* 2-pass, *TopHat2*, and the simulated ground truth alignment).

In general we find varying accuracies across the different event types, with consistent patterns for all the tested tools. Intron retentions are the most difficult to predict and exon skips the easiest. *rMATS* was able to detect only two kinds of events on the data we provided: exon skips and mutual exclusive exons. Only exon skips were part of our evaluation. All event types that were not predicted are shown as bars of height zero. We also would like to note, that the simulated data resembles a polyA selected library. When working with non-polyA selected, rRNA depleted libraries, performance will likely be worse, as incompletely spliced transcripts will be amongst the sequenced fragments, diluting the signal.

Across all event types, sample sizes and alignment methods *SplAdder* shows the best performance compared to the other tools. Although *rMATS* shows the highest precision on the predicted exon skip events (0.965, cf. Supplemental Fig. S-6), it has a considerably lower recall, thus affecting its overall performance. Further, it does not predict any of the other assessed types. In contrast *JuncBase* shows a generally high recall but predicts many false positive events, resulting in a low precision (cf. Supplemental Figs. S-6 and S-7).

A high read coverage has, in general, a positive effect on prediction accuracy with better results for the samples covered at a higher depth. However, we observed some instances where high coverage results in lower performance, most likely due to more false positives in the predicted set.

### 5.2 Event Quantification

For all events that were correctly predicted by each approach, we compared the associated PSI value to the ground truth computed on the simulated abundances.

In general, we observe good correlation between predicted and true PSI values (cf. Supplemental Table F for a list of all coefficients). Whereas *SplAdder* shows the highest correlation for exon skip events, *JuncBase* has slightly higher accuracy for the other event types, although closely followed by the SplAdder predictions. As *rMATS* only predicted exon skip events, we could only include this one event type into our comparison.

We did not observe large differences between correlation values for the different aligners. Interestingly, a higher read depth led to slightly lower quantification accuracies for all tools, even when using the unaligned ground truth read data. We speculate that this is an effect of the simulation tool. However, since we use the reads only for a relative comparison of the different approaches, our evaluation should not suffer from this.

### 5.3 Differential Analysis

*SplAdder* can be utilized in two different ways to compare alternative splicing between samples. One approach is to use the event mini-genes output by *SplAdder* as input to other tools for the analysis of differential transcript usage. For our experiments, we use *rDiff* and refer to this use case as *SplAdder*+*rDiff*. In addition, we recently added a testing module to the *SplAdder* core pipeline that uses a Generalized Linear Model (GLM), which we will refer to as *SplAdder*+*GLM* in the following evaluations. Based on the two artificial data sets described above, we find that *SplAdder* shows very good performance overall when compared to other testing approaches (Figure 3).

**Figure 3:**
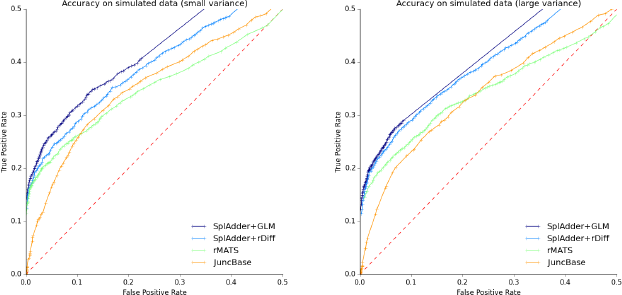
**Differential Testing Evaluation** Testing accuracy for four different methods (*SplAdder*+*GLM*, *SplAdder*+*rDiff*, *rMATS* and *JuncBase*; see legend). Each plot represents a different test set. The plot shown on the left represents the sample dataset with small biological variance between replicates, whereas the plot on the right is based on the sample set with increased biological variance between replicates. The dashed line represents the diagonal and reflects the performance of a random assignment of classes.

In the range of a low false positive rate, the performance of *SplAdder*+*rDiff* is comparable to *rMATS* and slightly inferior to *SplAdder*+*GLM*. This is consistent for both the small and large variance dataset. *JuncBase* uses a t-test for assessing the different groups of samples, which appears less well suited for testing read count data, as it leads to relatively many false positives at high confidence. The ROC curve shape directly reflects this.

### 5.4 Software and Usability

We have taken great care when implementing the *SplAdder* approach. It has been developed in Matlab but was translated into Python to improve accessibility. Both implementations provide the same functionality, however we will continue future development in Python only. When it comes to usability, *SplAdder* is a convenient one-stop-shop that provides all analysis within a single pipeline. With one simple command line call specifying the parameter set, all subsequent steps are automatized. In addition, the pipeline can be broken into single steps if necessary.

All other tested approaches required invocation of multiple separate tool components and required custom scripting on the user side to form a coherent pipeline. A single exception is *rMATS* that is also well engineered and is quite usable. Most of this also reflects in the running times of the implementations (cf. Supplemental Table E). Whereas *rMATS* and *SplAdder* have quite low running times, *JuncBase* and SpliceGrapher are considerably slower. Especially the *Cufflinks* preprocessing for *JuncBase* is very compute intense, with up to 30 hours for some evaluation samples of the largest size. Thus, we have excluded this preprocessing time from the running time table for *JuncBase*.

We believe that *SplAdder*’s improved usability is an important feature that will enable comprehensive AS analysis on RNA-Seq data for a wider audience than with previous methods. Our method is particularly timely, given the ubiquitous precence of available RNA-seq data, high interest in quantifying splicing phenotypes, and scalability to process thousands of samples.

## 6 Conclusion

We present *SplAdder*, a novel approach for the large-scale analysis of alternative splicing events based on RNA-Seq data. We also provide a thoroughly engineered software implementation that is straightforward to use and can be easily deployed in a high performance computing framework. *SplAdder* has been successfully applied to splicing analysis in various organisms, compares favorably to various other state of the art methods showing an overall high accuracy and can be readily applied to datasets of thousands of samples. We are working to further improve *SplAdder* to natively work with high performance compute clusters and generate more interactive visualizations.

## Acknowledgements

The authors are grateful to Vipin T Sreedharan for providing code to convert annotation files, to Andreas Wachter for valuable discussions and feedback on the software and to David Kuo for proofreading. *Funding* was provided by the Max Planck Society, Memorial Sloan Kettering Cancer Center, by the German Research Foundation (RA1894/2-1) and the Lucille Castori Center for Microbes, Inflammation, and Cancer (No. 223316).

